# Mathematical modeling of mechanosensing and contact-dependent motility coordination in *Myxococcus xanthus*

**DOI:** 10.1101/2022.12.02.518919

**Authors:** Yirui Chen, Jing Chen

## Abstract

Sensing and responding to mechanical cues in the environment are important for the survival and propagation of bacteria. A ‘social’ bacterium, *Myxococcus xanthus*, which exhibits periodic cell reversals, modulates its reversal frequency in response to environmental mechanical cues, such as substrate stiffness and cell-cell contact. In *M. xanthus* populations, the cell-cell contact-dependent reversal control is particularly important for formation of complex multicellular patterns and structures during the cooperative ‘social’ behaviors. Here we hypothesize that the gliding motility machinery of *M. xanthus* can sense the environmental mechanical cues during force generation and modulate the timing and frequency of cell reversal through signaling the cell’s reversal control pathway. To examine our hypothesis, we extend an existing mathematical model for periodic polarity switching (which mediates periodic cell reversal) in *M. xanthus*, and incorporate the experimentally suggested (i) intracellular dynamics of the gliding motors and (ii) interactions between the gliding motors and reversal regulators. The model results suggest the proper mode of interaction between the gliding motors and reversal regulators that can generate the observed increase of cell reversal frequency on stiffer substrates. Furthermore, the selected model predicts a cell reversal response to cell-cell contact, which is sufficient for generating the rippling wave, an important multicellular pattern in *M. xanthus* populations. Our model highlights a potential role of the gliding machinery of *M. xanthus* as a ‘mechanosensor’ that transduces mechanical cues into a reversal control signal.

## Introduction

The ability to sense the environment and adapt their behaviors accordingly is important for the survival and propagation of bacteria. For motile bacteria, their motility is often highly adjustable in response to environmental cues. The most well-studied example is chemotaxis in swimming bacteria like *E. coli*: the bacterium adjusts its frequency of random tumbling in response to the temporal change in the attractant or repellent concentration which it senses during swimming; in the long term, this amounts to biased movement towards attractants or away from repellents (Wadhams and Armitage, 2004;Sourjik and Wingreen, 2012). Besides diffusive chemical signals, bacteria can also adapt their motility (and other behaviors) to mechanical cues (Persat et al., 2015;Colin et al., 2019;Gordon and Wang, 2019;Dufrene and Persat, 2020). For instance, increasing load on the *E. coli* flagellum induces stabilization of the stator units in the flagellar motor, leading to recovery of the rotation frequency of the flagellum (Lele et al., 2013;Tipping et al., 2013;Wadhwa et al., 2019). For another example, Type IV pili, an appendage driving twitching motility in a broad range of bacteria, switches from extension to retraction rapidly and nearly exclusively upon touching a surface by its tip, suggesting its capability of sensing contacts (Tala et al., 2019;Dufrene and Persat, 2020). Because bacteria experience constantly fluctuating forces, especially when they live on inhomogeneous surfaces or in complex biofilms, their ability to respond to mechanical cues likely plays a critical role in helping them locate favorable niches and survive.

Although mechanosensing mechanisms have been intensively studied in eukaryotic cells both experimentally and theoretically (Cheng et al., 2017;Martino et al., 2018), the same topic is yet poorly investigated in bacteria. In this work, we focus on a cellular mechanism for mechanosensing in a soil-dwelling bacterium, *Myxococcus xanthus*, and its effect on the bacterium’s reversal frequency. *M. xanthus* moves on substrate surface and frequently reverses its direction of motion (Mauriello et al., 2010a;Bretl and Kirby, 2016;Schumacher and Sogaard-Andersen, 2017). Its reversal frequency varies under the influence of both chemical and mechanical cues. For the latter, physical contact with other *M. xanthus* cells (Shi et al., 1996;Welch and Kaiser, 2001) or prey cells (McBride and Zusman, 1996;Zhang et al., 2020), varying substrate stiffness (Zhou and Nan, 2017), etc., affect the reversal frequency of *M. xanthus* cells. Particularly, *M. xanthus* cells reverse on hard, 1.5% agar almost twice as frequently as they do on soft, 0.5% agar (Zhou and Nan, 2017). Modulation of cell reversal frequency is particularly crucial to emergence of various spatial patterns and structures in *M. xanthus* populations, such as streams, rippling waves, aggregation and fruiting bodies (Welch and Kaiser, 2001;Zhang et al., 2012a;Keane and Berleman, 2016). These spatial patterns and structures play important roles in “social” collaboration on feeding and sporulation in groups of *M. xanthus* cells (Berleman and Kirby, 2009;Velicer and Vos, 2009;Zhang et al., 2012a;Munoz-Dorado et al., 2016).

The molecular players underlying *M. xanthus* motility and reversal are mostly known. Force generation in *M. xanthus* is carried out by two motility systems: social (S-)motility favored by cells in large groups and adventurous (A-)motility favored by solitary cells (Mauriello et al., 2010a;Zhang et al., 2012a). The S-motility, also known as the twitching motility, is driven by Type IV pili (Sun et al., 2000;Mauriello et al., 2010a;Zhang et al., 2012a), the same cellular appendage driving twitching motility in *Pseudomonas aeruginosa* (Burrows, 2012). The A-motility, also known as the gliding motility, is powered by multi-subunit Agl-Glt complexes, which travel along helical intracellular trajectories and generate propulsion at the cell-substrate interface (Nan et al., 2011;Nan et al., 2013;Nan et al., 2014;Islam and Mignot, 2015;Faure et al., 2016). Both the S-and A-motility machineries are assembled or activated at the leading pole of the cell (Zusman et al., 2007;Mauriello et al., 2010a). The polarity of the cell is defined by asymmetric concentration of polarity-setting molecules at the two cell poles, particularly, MglA that concentrates at the leading pole, and MglB and RomR that concentrate at the trailing pole (Leonardy et al., 2007;Leonardy et al., 2010;Patryn et al., 2010;Zhang et al., 2010;Bulyha et al., 2011). Switching of the polarity-setting molecules between the two poles causes reversal of cell polarity and hence reversal of cell movement (Zhang et al., 2010;Zhang et al., 2012b;Treuner-Lange et al., 2015;Schumacher and Sogaard-Andersen, 2017;Guzzo et al., 2018;Szadkowski et al., 2019). Furthermore, the reversal frequency is controlled by the Frz signaling pathway, a chemosensory pathway with a cytoplasmic chemoreceptor, FrzCD (Astling et al., 2006;Zusman et al., 2007;Mignot and Kirby, 2008). Although the molecular mechanisms for force generation (Mignot et al., 2007;Nan et al., 2011;Sun et al., 2011;Nan et al., 2013;Faure et al., 2016) and polarity switching (Zhang et al., 2010;Zhang et al., 2012b;Treuner-Lange et al., 2015;Schumacher and Sogaard-Andersen, 2017;Guzzo et al., 2018;Szadkowski et al., 2019) have been quite intensively studied in *M. xanthus*, little is known about the mechanism regulating its reversal by mechanosensing.

A key component of a mechanosensing mechanism is molecules that transduce external mechanical cues into intracellular signals. One class of the most promising candidates for this role are motility molecules, which, by the nature of their function, form mechanical links between the cell and the external substrate to drive cell motility. In eukaryotes, for example, mechanosensing of the extracellular environment can indeed be mediated by molecules forming the focal adhesions (Cheng et al., 2017;Tao et al., 2017;Martino et al., 2018) – dynamic macromolecular structures that mechanically link the cell to the extracellular matrix and mediate cell migration. In *M. xanthus*, the A-motility machinery is connected to the molecules controlling cell polarity and cell reversal in several ways. First, the Agl-Glt machinery is activated by MglA and carries MglA as a subunit in its force-generating form (Patryn et al., 2010;Zhang et al., 2010;Treuner-Lange et al., 2015). Second, AglZ, another subunit of the Agl-Glt machinery, forms clusters anti-localized to the clusters of FrzCD (Mauriello et al., 2009;Nan et al., 2010).

These connections between the A-motility machinery and the polarity/reversal regulators suggest that the A-motility machinery could modulate cell reversal according to the mechanical cues it senses while propelling the cell.

To investigate the possibility that the A-motility machinery regulates cell reversal in *M. xanthus*, we extended an existing mathematical model for periodic polarity switching in *M. xanthus* (Guzzo et al., 2018) and incorporated (i) subcellular trafficking of the A-motility machineries and (ii) regulation of MglA and the Frz signal by the A-motility machinery. Our model shows that coupling between the A-motility machinery and MglA alone predicts a relationship between substrate stiffness and reversal frequency that is opposite to the experimental observation, but a selected mode of feedback from the A-motility machinery to the Frz signal can reverse the prediction and explain the experimental observation. We further used the model to investigate modulation of reversal frequency by physical contact between cells.

Exploiting a previous model for rippling wave formation in *M. xanthus* (Igoshin et al., 2001;Igoshin et al., 2004a), we were able to confirm that the reversal response predicted by our model can support emergence of rippling wave, an important multicellular pattern with strong implications in collaborative feeding (Berleman and Kirby, 2009;Keane and Berleman, 2016;Munoz-Dorado et al., 2016) and fruiting-body formation (Welch and Kaiser, 2001;Munoz-Dorado et al., 2016). Overall, our model proposes a feasible mechanism of mechanosensing in *M*.*xanthus*, which allows the bacterium to adapt its motility not only to its environment, but also to fulfill its “social” interaction with colony mates.

## Results

### Model with coupling between polarity control and A-motility

In light of the existing knowledge about the polarity switching mechanism and A-motility mechanism in myxobacteria, we constructed a model (**Figure 1**) combining the two mechanisms. The model is built upon the following key assumptions suggested by experimental observations. For short, in the remaining of the paper we will call the multi-subunit A-motility machinery as the “A-motor”.

1. An A-motor can assume three possible states: inactive, active or engaged. An inactive motor only diffuses in the cell, which likely represents disassembled parts of the motor (Nan et al., 2015;Faure et al., 2016;Nan, 2017;Wong et al., 2021). An active motor travels directionally along the helical track either towards the leading pole or towards the trailing pole. The directionality of an active motor randomly switches, as suggested by a previous experiment (Nan et al., 2015). (Because MreB filaments, the track that A-motor moves along (Mauriello et al., 2010b;Nan et al., 2011;Treuner-Lange et al., 2015;Fu et al., 2018), are short patches with indefinite polarity (Kim et al., 2006;Errington, 2015;Billaudeau et al., 2017), the motors may switch direction where the track switches polarity.) As an active A-motor passes through a focal adhesion site, it can engage with the focal adhesion and generate thrust on the cell. The engagement of a motor at the focal adhesion site likely represents dynamic complexing of the inner-membrane motor with the periplasmic and outer-membrane subunits, like engaging the clutch of a car.
2. Activation and deactivation of the A-motor occur at the cell poles, where the polarity setters are concentrated (Zhang et al., 2010;Zhang et al., 2012b;Treuner-Lange et al., 2015;Schumacher and Sogaard-Andersen, 2017;Guzzo et al., 2018;Szadkowski et al., 2019). The motor is activated via binding with MglA, which is reversed by MglB, as MglB inactivates MglA (Miertzschke et al., 2011;Keilberg and Sogaard-Andersen, 2014).
3. The dynamic localization of MglA/MglB/RomR at the two cell poles is described by a recent model proposed by Guzzo et al. (Guzzo et al., 2018). The model attributes polar oscillation of these molecules to feedback loops among them that regulate their binding to the poles. Specifically, MglA and MglB antagonize each other in polar binding; MglB promotes polar binding of RomR and its own polar binding; finally, RomR promotes polar binding of MglA. Strong mutual inhibition between MglA and MglB breaks the symmetry and concentrates them to opposite poles. The negative feedback loop, MglA --l MglB → RomR → MglA, causes periodic switching in their polar localization.

With the equations and parameters given in the **Supplementary Methods** (**Eqs. (S1)-(S39), Tables S1-S4**), our model recapitulates key motility characteristics observed experimentally, including:

1. The cell reverses periodically every ∼11 minutes (**Figure 2A**), consistent with the experimental data (Kaimer and Zusman, 2016;Guzzo et al., 2018;Szadkowski et al., 2019).
2. MglA is concentrated at the leading pole, and MglB and RomR are concentrated at the trailing pole (**Figure 2B**), agreeing with the experimental observations (Zhang et al., 2010;Zhang et al., 2012b;Treuner-Lange et al., 2015;Guzzo et al., 2018;Szadkowski et al., 2019).
3. In the cell body, MglA forms a concentration gradient decreasing from the leading pole to the trailing pole (**Figure 2D, E**). About half of the MglA are localized in the cell body (**Figure 2C**). The predicted spatial distribution of MglA is consistent with the experimental data (Nan et al., 2015).
4. Active motors also form a gradient that decreases from the leading pole to the trailing pole (**Figure 2D, E**), consistent with the observed distribution of AglZ clusters in elongated *M. xanthus* cells (Mignot et al., 2007).

**Figure 1:**
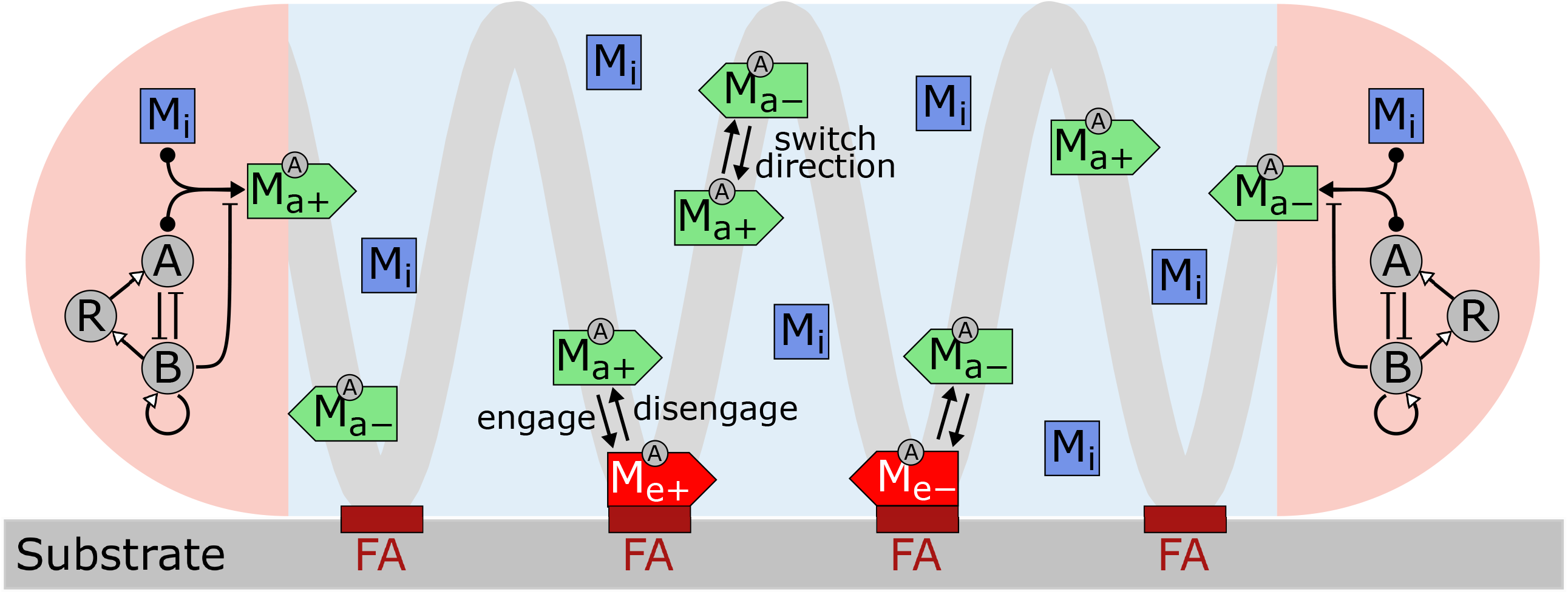
Model for *M. xanthus* reversal control with mechanosensing through A-motors. Motors are present in three states in the cell: inactive (M_i_), active (M_a+_, M_a−_) and engaged (M_e+_, M_e−_). Inactive motors diffuse. Active motors move directionally along the helical trajectory (light grey line) and randomly switch directions (+: moving to right; −: moving to left). When active motors pass a focal adhesion (FA) site (maroon bars), they can become engaged in force generation. Activation and deactivation of motors are confined within the cell poles (pink areas). Activation of a motor involves binding of MglA (A) and is reversed by MglB (B). Feedback loops among MglA (A), MglB (B) and RomR (R), proposed by Guzzo et al. (Guzzo et al., 2018), control periodic polarity switching. Arrows with solid head: state conversions/reactions. Arrows with solid heads and circle tails: reversible reactions. Arrows with hollow heads: positive regulations. Arrows with blunted heads: negative regulations. Regulations among MglA, MglB and RomR refer to regulations of their polar binding.

**Figure 2:**
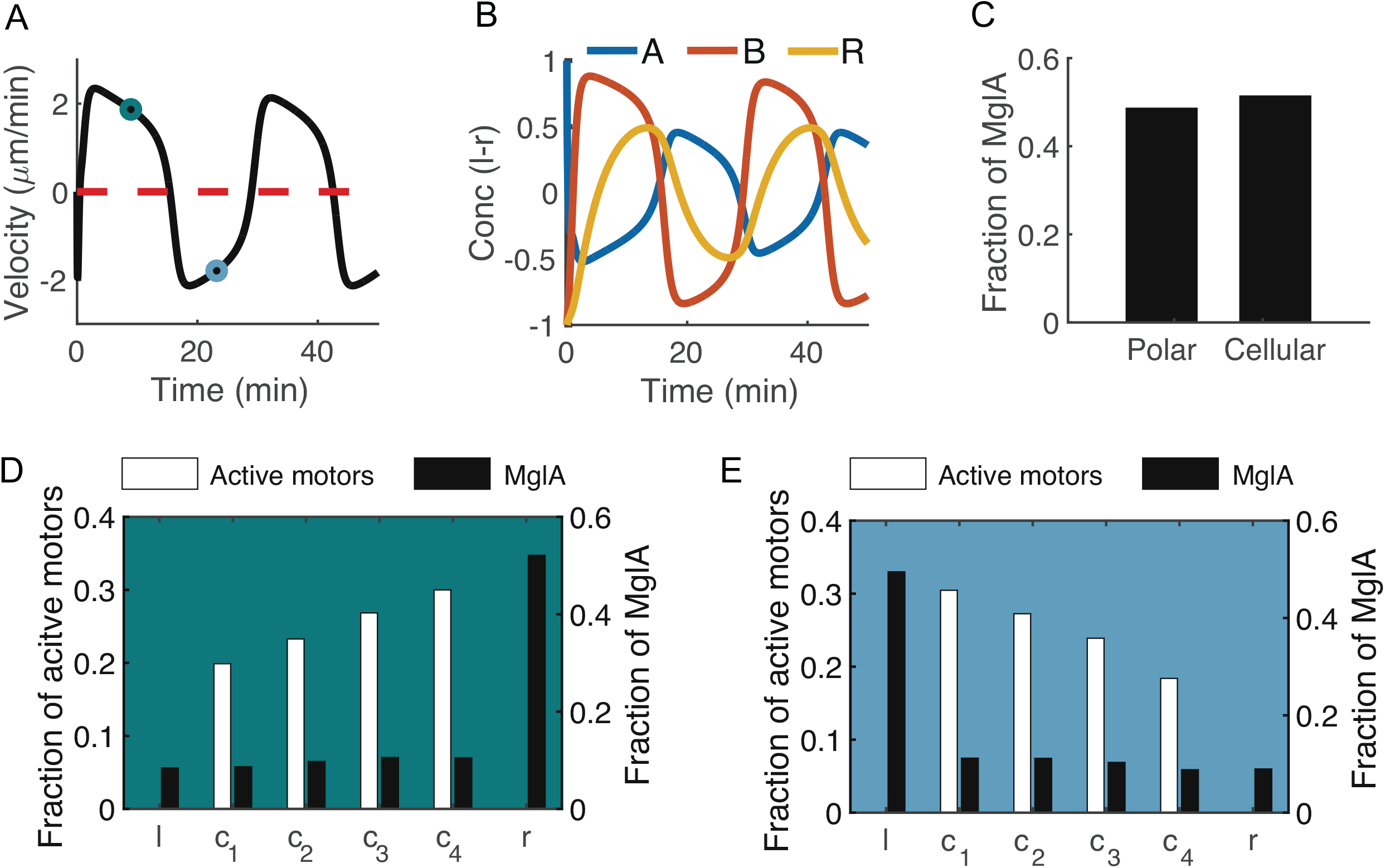
Key characteristics of *M. xanthus* motility predicted by the model. **(A**) Cell velocity over time. Positive velocity: right-moving. Negative velocity: left-moving. (**B**) Time courses of the concentration difference of MglA, MglB and RomR between the left and right poles. l: left pole. r: right pole. Positive values indicate concentration of the molecules at the left pole, and negative values indicate concentration at the right pole. (**C**) MglA distribution in the polar vs. cellular domains. (**D, E**) Spatial distribution of engaged motors and MglA along the cell when the cell is heading right (D) versus left (E). Black bars: MglA. White bars: engaged motors. l: left pole. r: right pole. c_1-4_: cellular subdomains. (D) and (E) correspond to the teal and blue time points, respectively, in (A).

### Interaction between Frz signaling and A-motors is necessary for explaining the response of cellular reversals to substrate stiffness

Because our model captures the physical interactions between the A-motors and the polarity setting molecule, MglA, the model can be used to predict the effect of external mechanical cues on the cell reversal frequency. We first examined if our model can explain the experimental observation that *M. xanthus* reverses more frequently on harder substrate surfaces, e.g., 1.5% agar versus 0.5% agar (Zhou and Nan, 2017). To characterize the impact of substrate stiffness on the A-motors, we note the previous observation that harder substrates intensify clustering of A-motors at the cell-substrate interface (Nan et al., 2010). Because clustered A-motors at the substrate interface are engaged in force generation (Mignot et al., 2007;Mauriello et al., 2010a;Sun et al., 2011;Nan et al., 2013), the phenomenon above indicates that harder substrates increase the number of engaged motors. In this light, we represented substrate stiffness by the motor engagement rate in the model: a harder substrate is represented by a higher engagement rate, and vice versa (**Figure 3C, D**). The value of the engagement rate was varied in a range such that the fraction of motile motors (i.e., summed fraction of active and inactive motors, or 1 – fraction of engaged motors) (**Figure 3C**) matches the experimental measurements on 0.5%-1.5% agar (Nan et al., 2013). We also varied the cellular drag coefficient between 30% and 200% of its default value to make the simulated cell velocity invariant to substrate stiffness, as observed in 0.5%-1.5% agar (Tchoufag et al., 2019); but this addition does not affect the predicted cell reversal frequency (**Figure S1**).

**Figure 3:**
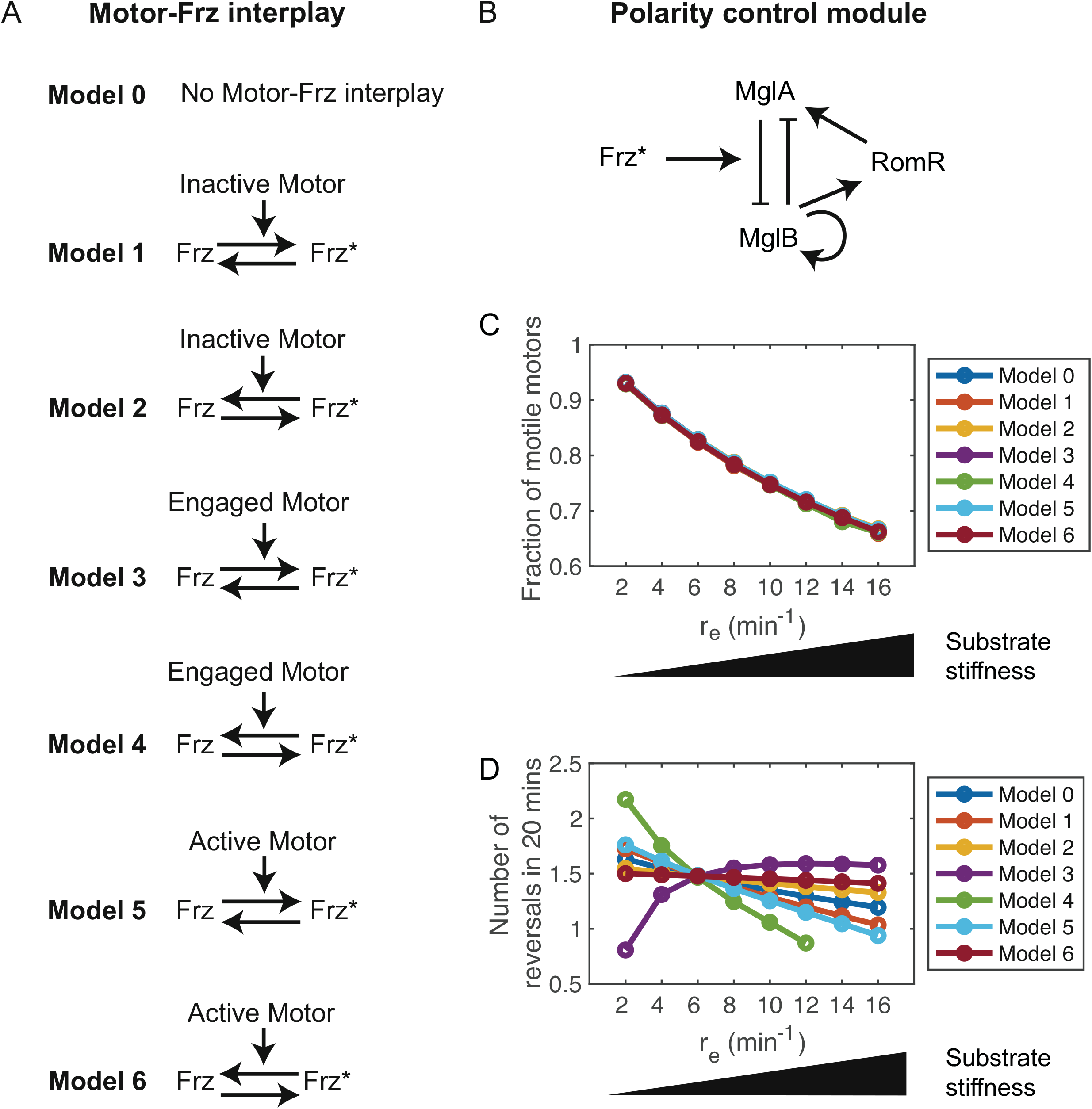
Impact of interplay between A-motor and Frz on the cellular response to varied substrate stiffness. (**A**) Possible modes of interplay between A-motors and Frz. Frz: inactive Frz. Frz*: active Frz. (**B**) Regulation of the MglA/MglB/RomR polarity control module by active Frz. Active Frz promotes release of MglB from the cell pole, following (Guzzo et al., 2018). This regulation is added to the polarity control module at the polar domain in Figure 1. (**C**) Predicted relationship between the motor engagement rate and fraction of motile motors in Models 0-6. Motile motors include inactive and active motors, but not engaged motors. (**D**) Predicted relationship between the motor engagement rate and cell reversal frequency in Models 0-6. The missing data points indicate that the model predicts no cell reversal.

With the substrate stiffness characterized by motor engagement rate, our model predicts a lower reversal frequency as the engagement rate increases, or as substrate stiffness increases (Model 0, **Figure 3**). To see why this is predicted by the model, recall that MglA binds with the active motors and engaged motors. An active motor can quickly reach either cell pole (∼2.5 s to traverse the typical cell length of 5 µm at an average speed of 2 µm/s (Nan et al., 2015)), and if it reaches the trailing pole, the motor can be inactivated and release MglA. In contrast, an engaged motor moves towards the trailing pole at a much slower velocity (equivalent to the cell speed, ∼2 µm/min, as the engaged motors are nearly stationary relative to the substrate (Mignot et al., 2007;Nan et al., 2010;Nan et al., 2013)). Therefore, engaged motors effectively sequester MglA in the cellular domain. Taken together, increasing the motor engagement rate boosts the number of engaged motors and hence sequesters MglA away from the polar domain. Note that MglA is a key component of the spatial oscillator, whose reduction at the poles inhibits polarity switching. This is how mechanical impacts on the motors affect cell reversal in the model. Of course, the prediction at this point contradicts the experimental observation (Zhou and Nan, 2017).

Next, we considered the effect of interaction between the A-motors and Frz signaling on the mechanosensing behavior in *M. xanthus*. Antagonistic spatial localization between FrzCD and A-motors reported previously (Mauriello et al., 2009;Nan et al., 2010) suggests potential interplay between FrzCD and A-motors. In this light, we investigated how such an interplay may further affect the cell reversal frequency in response to varied substrate stiffness, and whether adding this interplay makes the model prediction consistent with the experimental observation. FrzCD is the upstream protein of the Frz signaling pathway, which is known to interact with the MglA/MglB/RomR module and control its oscillatory frequency (Mignot and Kirby, 2008;Mercier and Mignot, 2016;Schumacher and Sogaard-Andersen, 2017). Hence, control of FrzCD by the A-motors, but not control of A-motors by FrzCD, can add to the regulation of reversal by A-motors in the cell. Since exactly how FrzCD is controlled by the A-motors is not known and many molecular details of the Frz pathway itself remain elusive (Mercier and Mignot, 2016), we built a collection of models representing all possible generic modes of control of the Frz signal by A-motors. Specifically, we assumed that activation or inactivation of the Frz signal can be promoted by A-motors in any of the three states: inactive, active, or engaged (**Figure 3A**). In our model, the Frz signal further regulates the MglA/MglB/RomR module in a similar way implemented by Guzzo et al. (Guzzo et al., 2018); particularly, high active Frz signal promotes release of MglB from the cell poles, which promotes polarity switching (**Figure 3B**).

Out of the six models with Frz-motor interaction, only Model 3 (engaged motors activate Frz) generates results that are opposite to Model 0 (without Frz-motor interaction) and qualitatively consistent with the experimental observations in (Zhou and Nan, 2017). In Model 3, increasing the engagement rate causes a strong increase of reversal frequency as substrate stiffness increases (**Figure 3D**); the change in reversal frequency is comparable to the experimental results (Zhou and Nan, 2017). Model 3 predicts an opposite dependency of the cell reversal frequency on substrate stiffness, because the increase of engaged motors in this case also promotes Frz activity. Higher Frz activity, in turn, promotes polarity switching in the model (Guzzo et al., 2018). The effect on Frz activity outcompetes the effect on MglA transport in Model 3, causing an overall increase of cell reversal frequency with increasing substrate stiffness. In summary, our model results suggest that interaction between A-motors and the Frz signal is essential for the observed mechanosensing behavior, and Frz is most likely activated by engaged motors.

### Our mechanosensing model explains contact-based reversal regulation

An important type of mechanical cue that regulates the cell reversal frequency in *M. xanthus* is physical contact between cells. Accelerated cell reversal induced by cell-cell contact is believed to be critical for formation of rippling waves (Igoshin et al., 2001;Igoshin et al., 2004a;Igoshin and Oster, 2004;Igoshin et al., 2004b;Sliusarenko et al., 2006), an important multicellular pattern with strong implications in collaborative feeding (Berleman and Kirby, 2009;Keane and Berleman, 2016;Munoz-Dorado et al., 2016) and fruiting-body formation (Welch and Kaiser, 2001;Munoz-Dorado et al., 2016). However, it remains unknown how physical contact between the cells transduces a signal to control cell reversal. Here we leveraged our model for *M. xanthus* mechanosensing to select possible mechanisms for the contact-based reversal regulation in *M. xanthus*. Based on the existing knowledge about the A-motility mechanism and surface properties of *M. xanthus*, we proposed two simplest mechanisms, by which cell-cell contact can affect A-motility and consequently affect reversal control.

A. The first hypothetical mechanism involves up-or down-regulating the engagement of A-motors in force generation. On the one hand (Hypothesis A1), cell-cell contact may increase the area of substrate surface (taking the neighboring cell as extra substrate surface) and promotes motor engagement. On the other hand (Hypothesis A2), if engagement of motors at the focal adhesion assumes strong cooperativity, increasing the substrate area may dilute the density of motors or motor-associated proteins located at the substrate interface, and hence reduce the effective rate of engagement.

B. The second hypothetical mechanism posits that the drag coefficient of the cell increases as it glides against another cell. Hypothesis B is based on the fact that myxobacterial cells are covered by fibrils and slime (Konovalova et al., 2010), which could cause a significant drag between two cells that slide against each other.

In the following subsections, we plugged each of the above hypothetical mechanisms into the mechanosensing model and examined the resulting reversal response to cell-cell contact. For better understanding of the model results, we will first present the results from the original model without Frz-motor interaction (Model 0), and then those from the selected model that matches the experimentally observed sensitivity to substrate stiffness (Model 3).

Firstly, we examined the cellular response generated by Hypothesis A using Model 0. As posited by the hypothesis, cell-cell contact transiently increases (Hypothesis A1, **Figure 4A, D**) or decreases (Hypothesis A2, **Figure 4I, L**) the motor engagement rate. The change was applied within a 3-minute time window to represent the typical time for cell-cell contact (time for two myxobacterial cells with a typical length of ∼6 µm to pass each other in opposite directions with a typical speed of ∼2 µm/min). The model results show that a transient increase in the motor engagement rate delays cell reversal and the level of reversal delay crucially depends on when the stimulus is applied during the reversal cycle (**Figure 4B, E, G**). If a transient increase in the motor engagement rate is applied shortly before a cell reversal, cell reversal is significantly delayed (**Figure 4B, G**). In contrast, the timing of cell reversal remains nearly unchanged when the stimulus is applied at the middle of the long steady phase between reversals (**Figure 4E, G**). This result can be understood via the help of phase portraits on the plane of the difference in MglA abundance between the two poles versus MglA abundance in the cellular domain (**Figure 4C, F**). Increase in the motor engagement rate induces an instant surge of MglA in the cellular domain (**Figure 4C, F**), because engaged motors sequester MglA in the cellular domain, as elaborated in the previous section. Since the stimulus is transient, the perturbation in MglA distribution is also transient. After the stimulus ends, the system quickly returns to the unperturbed state if it is yet far from cell reversal and proceeds as if it has never been perturbed (**Figure 4F**). If the perturbation ends close to cell reversal, however, the recent reduction in polar MglA would delay polarity switching and cell reversal (**Figure 4C**). To summarize, a transient increase in the motor engagement rate induces a phase-delay response with a sensitive period shortly before cell reversal and a refractory period elsewhere (**Figure 4H**). Conversely, if the stimulus is represented by a transient fall in the motor engagement rate, the model predicts a phase-advance response with similar sensitive and refractory periods (**Figure 4I-P**).

**Figure 4:**
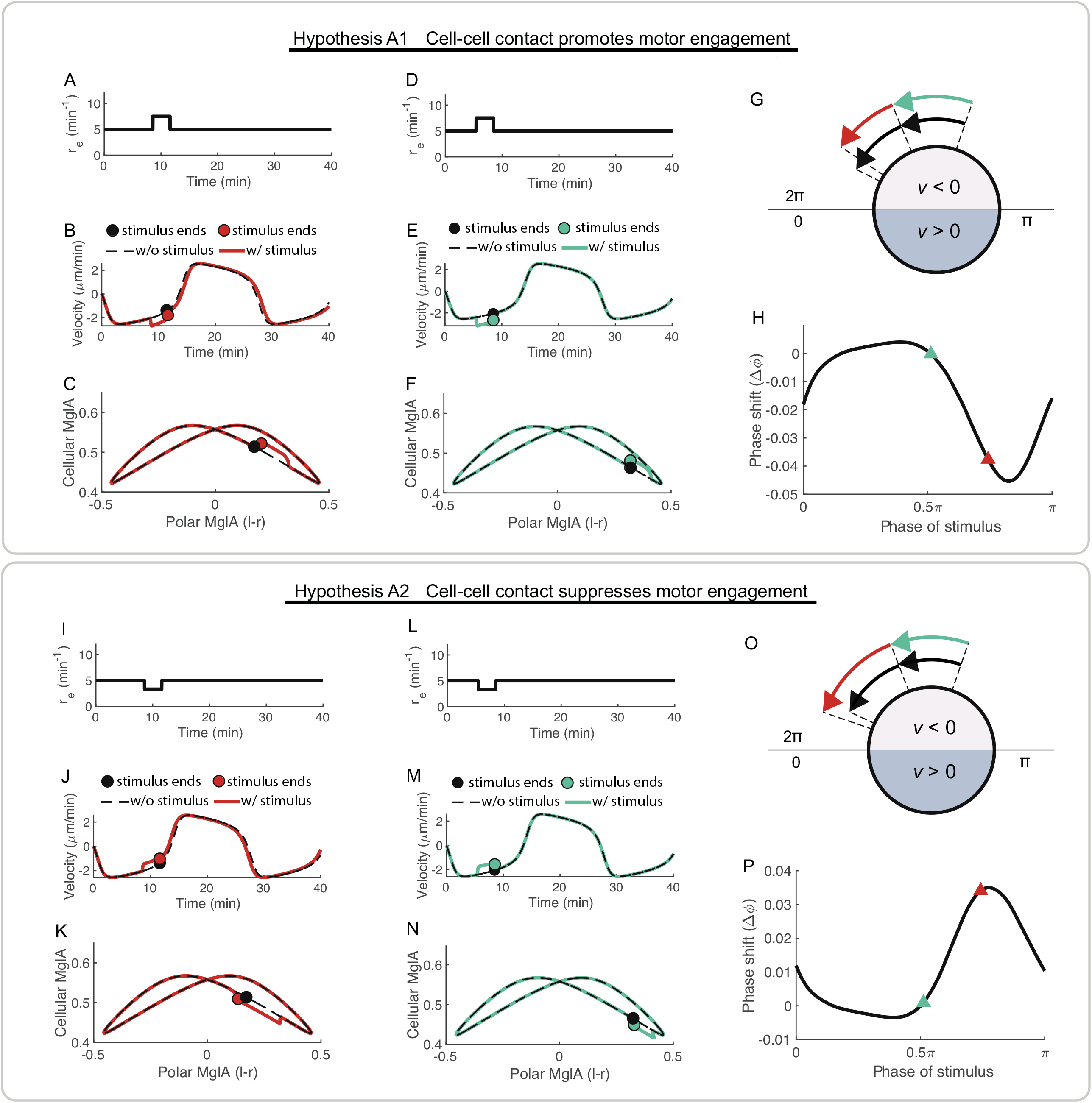
Contact-based reversal regulation by perturbation of motor engagement rate in the model without Frz-motor interaction. (**A-H**) Hypothesis A1: Cell-cell contact promotes motor engagement. The contact stimulus occurs in a 3-min time window right before a cell reversal (A-C) or during the steady phase between reversals (D-F). (**I-P**) Hypothesis A2: Cell-cell contact suppresses motor engagement. The contact stimulus occurs in a 3-min time window right before a cell reversal (I-K) or during the steady phase between reversals (L-N). (**A, D, I, L**) Time courses of the motor engagement rate representing the effect of cell-cell contact. (**B, E, J, M**) Simulated time trajectories of cell velocity with (solid lines) and without (dashed lines) the contact stimulus. The circles mark the end of the 3-minute stimulus window, and correspond to the dots with the same colors in (C, F, K, N). (**C, F, K, N**) Simulated trajectories on the phase plane of the difference of the fraction of MglA between the two poles and the fraction of MglA in the cellular domain. Same simulations as in (B, E, J, M). (**G, O**) Illustrative summary of the reversal responses to contact stimuli occurring at different times. Cell reversal is significantly changed if the contact stimulus occurs right before a cell reversal, but not if the stimulus occurs at the middle of the reversal cycle. Black arrows: progress of reversal cycle without contact stimulus. Red arrows: progress of reversal cycle with contact stimulus in (A) or (I). Teal arrows: progress of reversal cycle with contact stimulus in (D) or (L). (**H, P**) Predicted relationship between the phase shift in cell reversal and the timing of the contact stimulus. The phase shift is defined as Δ Δr · rr/T, where Δr is the shift in reversal time and *T* is the time interval between reversals without stimulus. The curve is same on the rr, 2rr) interval and not shown. Red triangles: data from (A, B) or (I, J). Teal triangles: data from (D, E) or (L, M).

Then we tested how transient changes in the motor engagement rate would affect cell reversal in the selected Model 3, where engaged motors activate the Frz signal. In this case, if a transient increase of the motor engagement rate (**Figure 5A**)is applied close to a cell reversal, the reversal is advanced (**Figure 5B, I**). Similar to the results from Model 0, a transient stimulus in the middle of the reversal cycle generates nearly no response (**Figure 5E, F, I**). Furthermore, a transient reduction in the engagement rate results in a phase delay with similar sensitive and refractory periods (**Figure 5K-T**). Compared to the phase response predicted by Model 0 (**Figure 4**), addition of the Frz-motor interaction reverses the phase response in Model 3 (**Figure 5**). Here is why. Like Model 0, a transient increase in the motor engagement rate in Model 3 boosts the number of engaged motors and hence moves some MglA from the polar domain to the cellular domain (**Figure 5C**). In Model 0, such a transient change in MglA distribution delays cell reversal (**Figure 4B, C**). In Model 3, however, the effect of MglA redistribution is outcompeted by the effect of concurring activation of the Frz signal by the increase of engaged motors (**Figure 5D**). Overall, Model 3 predicts a phase advance if cell-cell contact causes a transient increase in the motor engagement rate.

**Figure 5:**
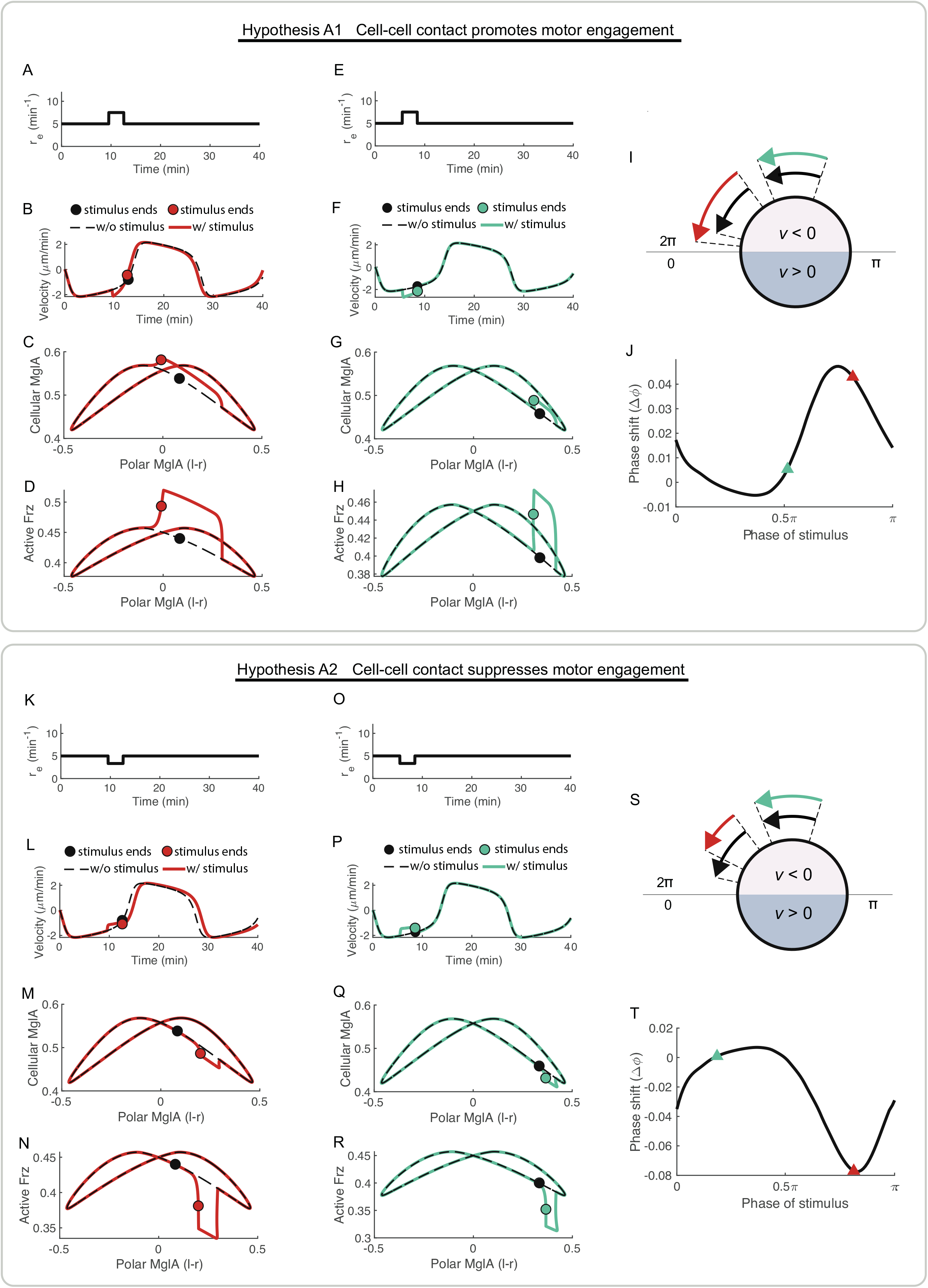
Contact-based reversal regulation by perturbation of motor engagement rate in the model with Frz-motor interaction. (**A-J**) Hypothesis A1: Cell-cell contact promotes motor engagement. The contact stimulus occurs in a 3-min time window right before a cell reversal (A-D) or during the steady phase between reversals (E-H). (**K-T**) Hypothesis A2: Cell-cell contact suppresses motor engagement. The contact stimulus occurs in a 3-min time window right before a cell reversal (K-N) or during the steady phase between reversals (O-R). (**A, E, K, O**) Time courses of the motor engagement rate representing the effect of cell-cell contact. (**B, F, L, P**) Simulated time trajectories of cell velocity with (solid lines) and without (dashed lines) the contact stimulus. The circles mark the end of the 3-minute stimulus window, and correspond to the dots with the same colors in (C, D, G, H, M, N, Q, R). (**C, G, M, Q**) Simulated trajectories on the phase plane of the difference of the fraction of MglA between the two poles and the fraction of MglA in the cellular domain. Same simulations as in (B, F, L, P). (**D, H, N, R**) Simulated trajectories on the phase plane of the difference of the fraction of MglA between the two poles and the activity of Frz. Same simulations as in (B, F, L, P). (**I, S**) Illustrative summary of the reversal responses to contact stimuli occurring at different times. Cell reversal is significantly changed if the contact stimulus occurs right before a cell reversal, but not if the stimulus occurs at the middle of the reversal cycle. Black arrows: progress of reversal cycle without contact stimulus. Red arrows: progress of reversal cycle with contact stimulus in (A) or (K). Teal arrows: progress of reversal cycle with contact stimulus in (E) or (O). (**J, T**) Predicted relationship between the phase shift in cell reversal and the timing of the contact stimulus. The phase shift is defined as Δ Δr · rr/T, where Δr is the shift in reversal time and *T* is the time interval between reversals without stimulus. The curve is same on the rr, 2rr) interval and not shown. Red triangles: data from (A, B) or (K, L). Teal triangles: data from (E, F) or (O, P).

Finally, we examined the reversal response under Hypothesis B using Model 0 and Model 3. In this case, the drag coefficient of the cell is transiently increased within a 3-minute time window to represent the contact stimulus (**Figure S2A, C, F, H**). In both Model 0 and Model 3, such a transient stimulus delays cell reversal slightly if it occurs shortly before cell reversal, but not other times during reversal cycle (**Figure S2E, J**). Cell reversal is delayed in this case because increasing the drag coefficient lowers the cell speed. Since the engaged motors carry MglA and travel at the cell speed towards the trailing pole of the cell (engaged motors are stationary with respect to the substrate (Mignot et al., 2007;Nan et al., 2010;Nan et al., 2013)), lowering the cell speed essentially reduces the flux of MglA into the trailing pole and hence delays polarity switching. However, this effect is weak and the shift of reversal time caused by changing the cellular drag coefficient is much less than that caused by a similar level of change in the motor engagement rate (compare **Figures S2** with **Figures 4, 5**).

In summary, our model results show that cell-cell contact is likely to modulate cell reversal frequency mainly through its effect on engagement of A-motors at the focal adhesions. Particularly, our selected Model 3 predicts a phase advance if cell-cell contact induces a transient increase in the motor engagement rate, and vice versa.

### Predicted reversal response can support rippling wave formation in M. xanthus population

A previous model by Igoshin et al. (Igoshin et al., 2001;Igoshin et al., 2004a) shows that a rippling wave like those observed in experiments can form in a *M. xanthus* population if cell-cell contact induces sufficiently strong phase advance with a refractory period right after cell reversals. This condition is qualitatively consistent with the reversal response predicted by our selected mechanosensing model (Model 3) when the contact stimulus is represented by a transient increase in the motor engagement rate (Hypothesis A1, **Figure 5A-J**). To quantitatively test if the reversal response predicted by Model 3 is sufficient for generating rippling waves in a *M. xanthus* population, we plugged the predicted response into Igoshin’s model and examined the resulting spatiotemporal patterns (**Figure 6A**).

**Figure 6:**
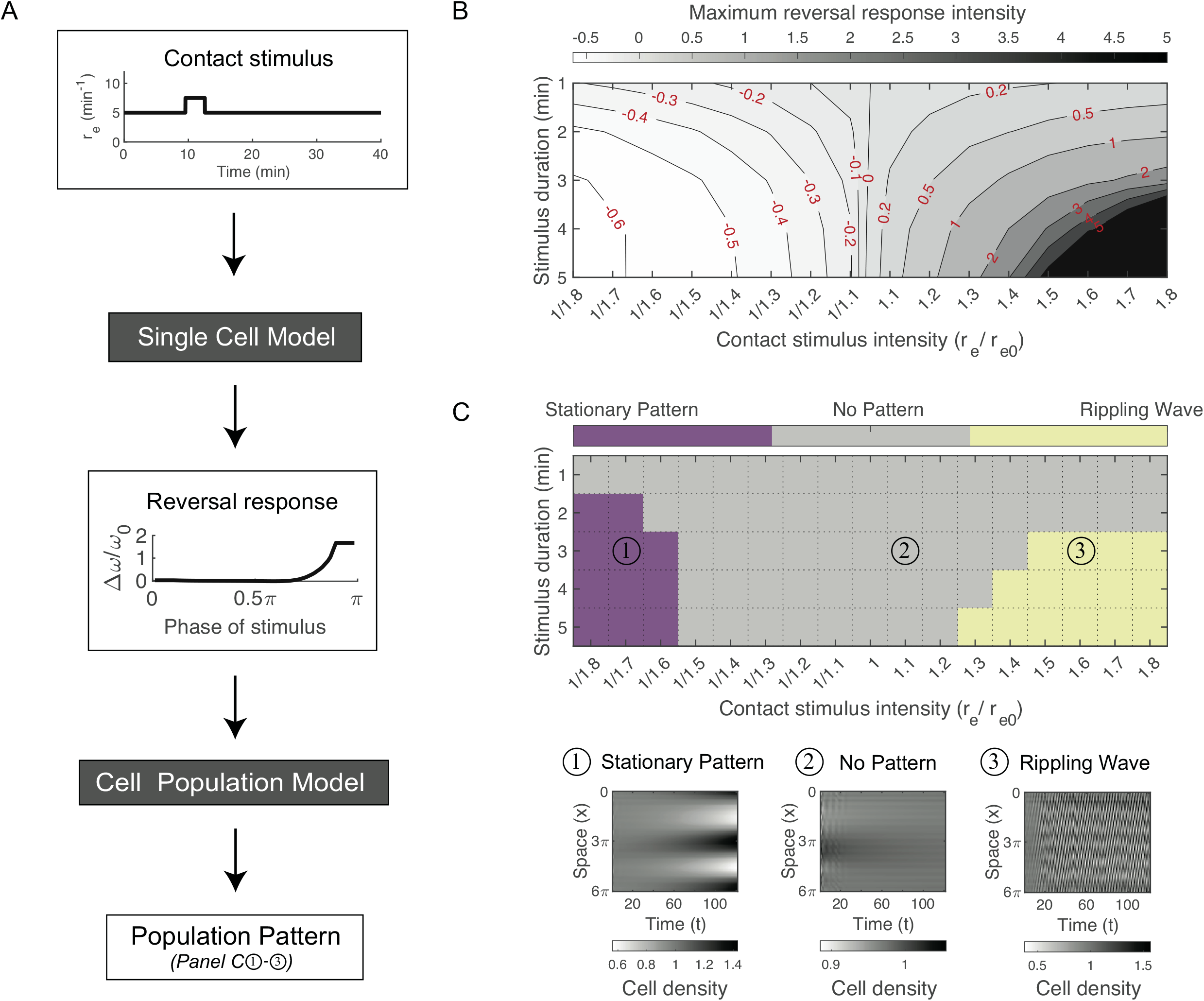
Predicted phase response can support rippling wave formation. (**A**) Workflow for modeling population pattern formation based on predicted phase response from the single-cell model. For contact stimuli imposed at different phases over the reversal cycle (e.g., **Figure 5A, E**), the single-cell model predicts a phase response curve (e.g., **Figure S3B, D**). The phase response curve is then plugged into Igoshin’s population model to predict the resulting population pattern. (**B**) Maximum phase response as a function of intensity and duration of the stimulus predicted from the single-cell model. Maximum phase response refers to the Δw with the highest absolute value on a phase response curve. The contact stimuli are represented by a transient increase or decrease in the motor engagement rate. (**C**) Population patterns generated with different stimulus durations and intensities. Bottom panel shows example patterns generated by the labeled parameter sets in the upper panel. The time (x-axis) and spatial location (y-axis) are non-dimensionalized (see **Supplementary Methods**).

As a mean-field model, Igoshin’s model (**Supplementary Methods, Eqs. (S40)-(S46)**) does not resolve individual cells, but rather describes the mean cell density as a variable dependent on time, space and phase in the reversal cycle. The cell density variable undergoes advection both in space and in the reversal phase (**Eq. (S40)**). The reversal response to cell-cell contacts is expressed in the model as a positive dependence of the phase velocity on the summed local density of cells moving in the opposite direction (**Eqs. (S42)-(S43)**). The phase velocity also depends on the phase itself to represent sensitive and refractory periods of the reversal response (**Eq. (S41)**). To utilize Igoshin’s rippling wave model, therefore, we need to convert Δ ϕ, the phase shift predicted by our single-cell model, into a change in the phase velocity.

To make the conversion, note that the default phase velocity without the stimulus is ω_0_ = π/reversal time. Assume that a stimulus occurs at a phase, 0 ≤ *ϕ*._s_ < *π*. Without the stimulus the cell would reverse in time, (*π*;-*ϕ*_s_)/*ω*_0_. With the stimulus the reversal occurs in time,

(*π*-*ϕ*_s_)/ *ω*_0_ -Δτ, where Δτ is the shift of reversal time and is related to Δ *ϕ*by Δ *ϕ* =_0_ Δτ. The effective phase velocity after the stimulus can hence be inferred from the ratio of the two waiting times for reversal, i.e.,

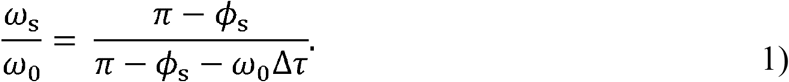

Then, the change in phase velocity, Δ*ω* = *ω*_s_ -*ω*_0_, which enters Igoshin’s population dynamics model, is written as Eq. (2). Note that Δ*ϕ*, Δτ and Δ*ω* all depend on *ϕ*_S_.

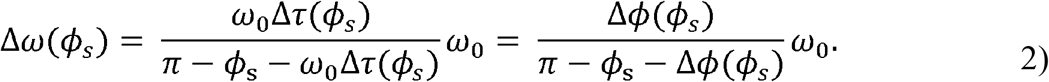

An advanced reversal with Δτ > 0 corresponds to an increase in the phase velocity, Δ*ω* > 0, and vice versa for a delayed reversal. By symmetry the same conversion applies to *π* ≤. *ϕ*_S_. < 2*π*, where the phase is folded to the [0, *π*] a range by a 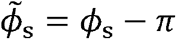 and 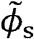a replaces*ϕ*_S_in **Eq. (2)**.

Using Eq. (2), we converted the phase shift (**Figure 5J, T**) predicted by our single-cell model into a phase response curve that Igoshin’s model adopts (**Figure S3**). Note that the phase response curves show consistent sensitive and refractory periods for different durations of stimuli if the phase of stimulus is accounted for by when it ends (**Figure S3B, D**), not when it begins (**Figure S3A, C**). This is consistent with our understanding of the reversal response elaborated above: whether the timing of cell reversal is affected mainly depends on if the system can return to the steady phase between reversals after the transient stimulus ends. Thereby, for the rest of the paper, the phase of stimulus refers to the ending phase of the stimulus. Note that a bacterium may contact more than one colony mate during a reversal cycle or pass a colony mate only half-way before both cells reverse; therefore, we need to consider stimuli with different durations and intensities. Overall, **Figure 6B** shows that the intensity of the reversal response increases as either the intensity or duration of the stimulus increases.

When we plugged the phase response predicted by our single-cell model into Igoshin’s model (Igoshin et al., 2001), we found that the rippling wave pattern only emerges with the stimulus represented as a transient rise in the motor engagement rate (i.e., Hypothesis A1) and requires sufficient intensity and/or duration of the stimulus (**Figure 6C**). In this case, the stimulus induces a phase advance with a refractory period during the early part of the reversal cycle (**Figure 5J**). Although a transient fall in the motor engagement rate can also provide a strong response as a phase delay (**Figure 6B**), the phase delay only gives rise to a stationary pattern, instead of a rippling wave (**Figure 6C**). In summary, our finding is consistent with Igoshin’s conclusion that rippling wave emerges with sufficient phase advance with a refractory period right after cell reversal, and the phase response predicted by our single-cell mechanosensing model with a contact-induced transient increase in the motor engagement rate is sufficient for generating the rippling wave.

## Discussion

In this work, we developed the first mathematical model for mechanosensing-based reversal control in *M. xanthus*. The model highlights the interplay between the reversal control pathway and A-motors in *M. xanthus* mechanosensing (**Figure 1**). The model not only reproduced the key characteristics of *M. xanthus* motility and reversal (**Figure 2**), but also proposed explanation for (1) how *M. xanthus* cells adapt their reversal frequency to varied substrate stiffness (**Figure 3**) and (2) how this mechanosensing mechanism allows *M. xanthus* cells to modulate their reversal upon physical contact with colony mates (**Figures 4 and 5**). We found that interplay between the A-motors and the Frz signal is necessary to reproduce the experimentally observed sensitivity of the cell reversal frequency to the substrate stiffness.

Particularly, activation of Frz by motors engaged in force generation (Model 3) is the most promising mode of Frz-motor interplay (**Figure 3**). Moreover, if cell-cell contact induces a transient increase in the motor engagement rate, Model 3 predicts a phase advance in cell reversal that is sufficient to generate rippling waves in a *M. xanthus* population (**Figure 6**). Our model proposes that the A-motility machinery of *M. xanthus* potentially serves as a ‘mechanosensor’ and transduces mechanical cues in the environment into a signal modulating cell reversal.

We note that our model predictions are semi-quantitative, because many quantitative details about the reversal control pathway and A-motility machinery remain elusive. For example, the interaction between the Frz proteins and A-motility machineries is suggested by the antagonistic spatial distribution between the force-generating motor clusters and the upstream receptor of the Frz pathway, FrzCD (Mauriello et al., 2009;Nan et al., 2010); but the exact interaction is unknown. For another example, the mechanistic details of force generation by the A-motility machineries are yet elusive, and hence it remains unclear how the machinery mechanically interact with the substrate, and how it mechanically interacts with a contacting cell. In the model, we resorted to a generic and simplistic assumptions that the motor engagement rate depends on substrate stiffness and cell-cell contact, based on the observed relationship between the motor clustering intensity and substrate stiffness (Nan et al., 2010). Nevertheless, by taking maximum usage of the existing information, our model is able to suggest what is necessary for the observed cellular behaviors in *M. xanthus*. Specifically, the model results demonstrate the necessity of Frz-motor interaction and suggest a transient increase in the motor engagement rate as the likely effect of cell-cell contact. These proposals point out interesting directions for future experimental investigation, such as dissecting the interaction between Frz proteins and the A-motility machinery, and tracking the clustering of A-motility machineries upon cell-cell contact. Results from these future experiments will also provide useful information for validation and revision of our mechanosensing model.

An important point to note is that the conclusion of Igoshin’s population model depends on the location of the refractory period in the reversal cycle. Our current mechanosensing model predicts a phase response with a refractory period during the early part of the reversal cycle, i.e., a period right after reversal. In this case, a phase advance is necessary to generate a rippling wave, whereas a phase delay would generate a stationary pattern (**Figures 6C**). However, if the refractory period is located during the late part of the reversal cycle, rippling wave would emerge in the case of phase delay, instead of phase advance (**Figure S4**). Note, the polar distribution of the polarity setting molecules in *M. xanthus* exhibits a long, nearly steady, phase between cell reversals, and switches swiftly right before reversals. Such dynamics indicates that the core part of the system works like a toggle switch with two symmetric steady states. Polarity switching is likely triggered by a component which slowly accumulates between reversals and quickly reset after a reversal, such as the polar distribution of RomR suggested by Guzzo et al. (Guzzo et al., 2018). If such a mechanism is confirmed, the system is indeed more sensitive to external perturbations at times closer to cell reversal – because at such times the level of the trigger component is close to the trigger threshold. Nevertheless, it is interesting for future studies to find out if some control mechanism may generate a refractory period during the late part of the reversal cycle. If *M. xanthus* reversal control works in this alternative manner, the conclusions of this work should be reversed.

Ultimately, *M. xanthus* motility control is probably influenced by a combination of mechanical and chemical stimuli. The Frz pathway that controls the cell reversal frequency is one of the eight chemotaxis-like pathways in *M. xanthus* (Zusman et al., 2007); FrzCD itself is homologous to the chemoreceptors. Moreover, the Frz pathway crosstalks with other chemosensory pathways, such as the Dif pathway (Xu et al., 2008), which senses lipids and controls *M. xanthus* motility as well (Bonner et al., 2005;Xu et al., 2011). It is not surprising that *M. xanthus* exploits multiple sensing mechanisms to orchestrate formation of numerous types of population patterns at different stages of its social life cycle (Keane and Berleman, 2016). It would be interesting for future studies to elucidate how mechanical and chemical stimuli jointly modulate the motility of *M. xanthus* to achieve the diverse population patterns.

Mechanosensing is likely an important function for bacterial cells, which allows the cell to ‘perceive’ the properties of the surfaces it gets into contact with and make beneficial decisions. Due to the difficulty of simultaneously tracking mechanic forces and signaling activities in a cell, mathematical modeling provides a powerful tool to integrate the two aspects of the dynamics under coherent mechanistic frameworks, and generate useful insights and/or predictions to facilitate experimental investigation. Ultimately, the integration of modeling and experimentation will provide the best tool to uncover mysteries in bacterial mechanosensing.

## Acknowledgements

This work was supported by NIH (1R35GM138370) and startup funds from the Virginia Tech Department of Biological Sciences and College of Science to JC. We further acknowledge members of the Chen lab for helpful discussion.

